# Chromatin accessibility and microRNA expression in nephron progenitor cells during kidney development

**DOI:** 10.1101/2021.03.05.434138

**Authors:** Andrew Clugston, Andrew Bodnar, Débora Malta Cerqueira, Yu Leng Phua, Alyssa Lawler, Kristy Boggs, Andreas Pfenning, Jacqueline Ho, Dennis Kostka

## Abstract

Mammalian nephrons originate from a population of nephron progenitor cells (NPCs), and it is known that NPCs’ transcriptomes change throughout nephrogenesis during healthy kidney development. To characterize chromatin accessibility and microRNA (miRNA) expression throughout this process, we collected NPCs from mouse kidneys at embryonic day 14.5 (E14.5) and postnatal day zero (P0) and assayed cells for transposase-accessible chromatin and small RNA expression. We observe 46,374 genomic regions of accessible chromatin, with 2,103 showing significant changes in accessibility between E14.5 and P0. In addition, we detect 1,104 known microRNAs, with 114 showing significant changes in expression. Genome-wide, changes in DNA accessibility and microRNA expression highlight biological processes like cellular differentiation, cell migration, extracellular matrix interactions, and developmental signaling pathways such as Notch. Furthermore, our data identify novel candidate cis-regulatory elements for *Eya1* and *Pax8*, both genes with a role in NPC differentiation; we also associate expression-changing microRNAs, including *let-7-5p*, *miR-125b-5p*, *miR-181a-2-3p*, and *miR-9-3p,* with candidate cis-regulatory elements. Overall, our data characterize NPCs during kidney development and point out new candidate regulatory elements for genes and microRNA with key roles in nephrogenesis.

## Introduction

The functional unit of the kidney is the nephron, which serves to filter waste and maintain normal homeostasis of water, acid-base, and electrolytes in the body. Nephron number varies widely in humans (typically between two hundred thousand and greater than one million nephrons (Hughson et al., 2003)) and is established prior to birth (birth in humans, approximately post-natal day 2-3 in mice (Hartman et al., 2007; Hinchliffe et al., 1991)). Nephrons cannot regenerate after birth, and decreased nephron endowment is associated with an increased risk of chronic kidney disease and hypertension (Bertram et al., 2011). Nephron number, in turn, is in large part determined by a population of cells called nephron progenitor cells (Cebrian et al., 2014a). During kidney development, one subset of nephron progenitor cells undergo mesenchymal-to-epithelial transitions (MET) to differentiate into cells of the early developing nephron (renal vesicle), while another continues to self-renew. In the latter stages of embryonic development, nephron progenitors increase their propensity to differentiate, which gradually depletes their population and marks the eventual cessation of nephrogenesis (Hartman et al., 2007; Rumballe et al., 2011; Volovelsky et al., 2018).

Nephron progenitor differentiation is regulated by a series of coordinated events. Briefly, Bmp7-pSmad1/5/8 signaling induces the initial exit of self-renewing Cited1^+^/Six2^+^ nephron progenitors into a primed Cited1^-^/Six2^+^ state, followed by Wnt9b/β-catenin induction of differentiation (McMahon et al., 2016; Rumballe et al., 2011). Migration of nephron progenitors around the ureteric bud influences their exposure to ureteric bud-secreted Wnt9b, prolonged exposure to which results in the differentiation of progenitors into *Wnt4*-expressing renal vesicles (Lawlor et al., 2019; Stark et al., 1994). Transcriptional changes in nephron progenitors occur as kidney development proceeds, and they are associated with genes and pathways that influence stem cell aging, in particular *mTor* and its repressor *hamartin* (Chen et al., 2015; Volovelsky et al., 2018). These changes are thought to desensitize nephron progenitors to signals for self-renewal, and in this way contribute to their depletion and the consequent cessation of nephrogenesis (Chen et al., 2015).

Recent studies have implicated non-coding RNA, such as microRNA (miRNA), to play a role in the regulation of nephrogenesis. miRNA are short, non-coding RNA molecules that target messenger-RNA transcripts (mRNA) for decreased translation or enhanced degradation, in both cases via the RNA-induced silencing complex (RISC) (Bartel, 2009). Loss of miRNA processing in nephron progenitors results in premature depletion of the nephron progenitor population (Ho et al., 2011); for instance, global removal of the hypoxia-responsive *miR-210* causes a significant decrease in nephron endowment in male mice (Hemker et al., 2020), and deletion of the *miR-17∼92* cluster in nephron progenitors impairs progenitor proliferation and reduces nephron endowment (Marrone et al., 2014). Further, the protein Lin28b is a known repressor of the *let-7* family of miRNAs (Heo et al., 2008), and its expression in nephron progenitors decreases as nephrogenesis proceeds. This reduction coincides with an increase in expression of the *let-7* family, and ectopic treatment with *Lin28b* is sufficient to prolong nephrogenesis (Yermalovich et al., 2019). This suggests that miRNA in the *let-7* family play an important role in timing the cessation of nephrogenesis.

In addition to transcriptional changes, changes in chromatin accessibility have been observed in nephron progenitor cells during embryogenesis, suggesting a developmentally-timed opening or closing of gene regulatory sequences, like promoters and enhancers (Hilliard et al., 2019). Consistent with this idea, transcription factors that regulate nephron progenitors including Six2, Hoxd11, Osr1, and Wt1 have been shown to bind enhancer sequences located in “regulatory hot-spots” in the genome (O’Brien et al., 2018), and the multipotent nephron progenitor marker Six2 co-binds enhancers with β-catenin to drive expression of marker genes of self-renewal and differentiation (Park et al., 2012).

In this study, we sought to identify enhancers and miRNAs in nephron progenitors that contribute to the cessation of nephrogenesis. While enhancers that regulate long non-coding RNA expression during nephrogenesis have been identified (Nishikawa et al., 2019), none have been shown to regulate miRNA expression in nephron progenitors. Therefore, to observe changes in miRNA expression and chromatin accessibility between embryonic day 14.5 (E14.5) and post-natal day zero (P0), we produced matched smRNA-seq and ATAC-seq datasets from nephron progenitor cells at these two time points. This allowed us to identify 2,103 regions of changing chromatin accessibility, and 114 miRNAs that are differentially expressed in nephron progenitor cells between these two time points. Enrichment analyses of chromatin regions with changing accessibility or among the predicted gene targets of differentially expressed miRNAs implicate pathways affecting epithelial cell differentiation, cell migration, extracellular matrix interactions, and key developmental signaling pathways such as Wnt, Notch, and TGF-β signaling. We observe significant changes in accessibility in chromatin, and present multiple lines of evidence implicating two genomic regions as (predicted) novel enhancers for the genes *Eya1* and *Pax8* in nephron progenitor cells. Finally, we identified consistent accessibility and miRNA expression changes for 33 pairs of miRNAs and accessible regions, suggesting gene regulatory loci for the expression of for miRNAs with a role in nephrogenesis, like *let-7-5p*, *miR-125b-5p*, *miR-181a-2-3p*, and *miR-9-3p*.

## Results

### Isolation of nephron progenitor cell populations from E14.5 and P0 kidneys

NP cells were isolated by positive selection for Itgα8 at E14.5 or at P0, and each sample was divided to perform ATAC-seq, smRNA-seq, and quality control **(Supplemental figure 1A-B**). Quantitative PCR (qRT-PCR) confirmed that the NP marker genes *Six2* and *Cited1* (Huang et al., 2016) are highly expressed in isolated nephron progenitor samples relative to whole kidney, more so than markers for other cell types (including ureteric bud (*Calb)* (Cebrian et al., 2014b), endothelial cells (*Pecam*) (Kobayashi et al., 2008), renal stroma (*Pdgfrβ*) (Chen et al., 2015) and renal vesicle (*Lhx1)* (Brunskill et al., 2014) (**Supplemental figure 1C**). With regard to sex, litters used for E14.5 samples ranged from 27% to 50% female, and litters for P0 samples ranged from 31% to 75% female. On average, E14.5 samples were 38% female, and P0 samples were 49% female **(Supplemental figure 1D**). These carefully age-matched NP cell populations allowed us to analyze chromatin accessibility and small RNA expression in the same stages of development at two time points (E14.5 and P0).

### Early and late nephron progenitors have distinct chromatin environments

ATAC-seq data from E14.5 and P0 nephron progenitors allowed us to identify 46,374 regions of accessible chromatin (with an irreducible discovery rate (Boleu et al., 2015) threshold of 0.1, see **Supplemental figure 2C, Supplemental table 1**), combined across samples and time points. We note that our data conformes to the ENCODE project’s ATAC-seq guidelines (ENCODE, n.d.) (**Supplemental figure 2A-B**), and that we observe increased rates of conservation among IDR peaks than are seen in randomized controls (**Supplemental figure 2D**). Among the IDRs we report, 3,576 contain enhancers documented in the VISTA (Visel et al., 2007) and / or FANTOM5 (De Rie et al., 2017) enhancer databases, and 13,495 contain DNA sequences with predicted regulatory roles in at least one mouse tissue or cell type according to EnhancerAtlas 2.0 (Gao and Qian, 2020).

Principal component analysis (adjusted for sex differences between samples) revealed a clear grouping of samples by developmental time point (**Figure 1A**). To identify genomic loci with changes in chromatin accessibility, we compared ATAC-seq signal between E14.5 and P0 samples. We observed 2,103 differentially accessible regions (DARs), controlling the false discovery rate at 10%. Of these 1,323 (55%) showed increased accessibility at P0 compared with E14.5, and hierarchical clustering revealed a clear distinction between chromatin regions that open and close between E14.5 and P0, respectively (**Figure 1B**).

**Figure 1.**
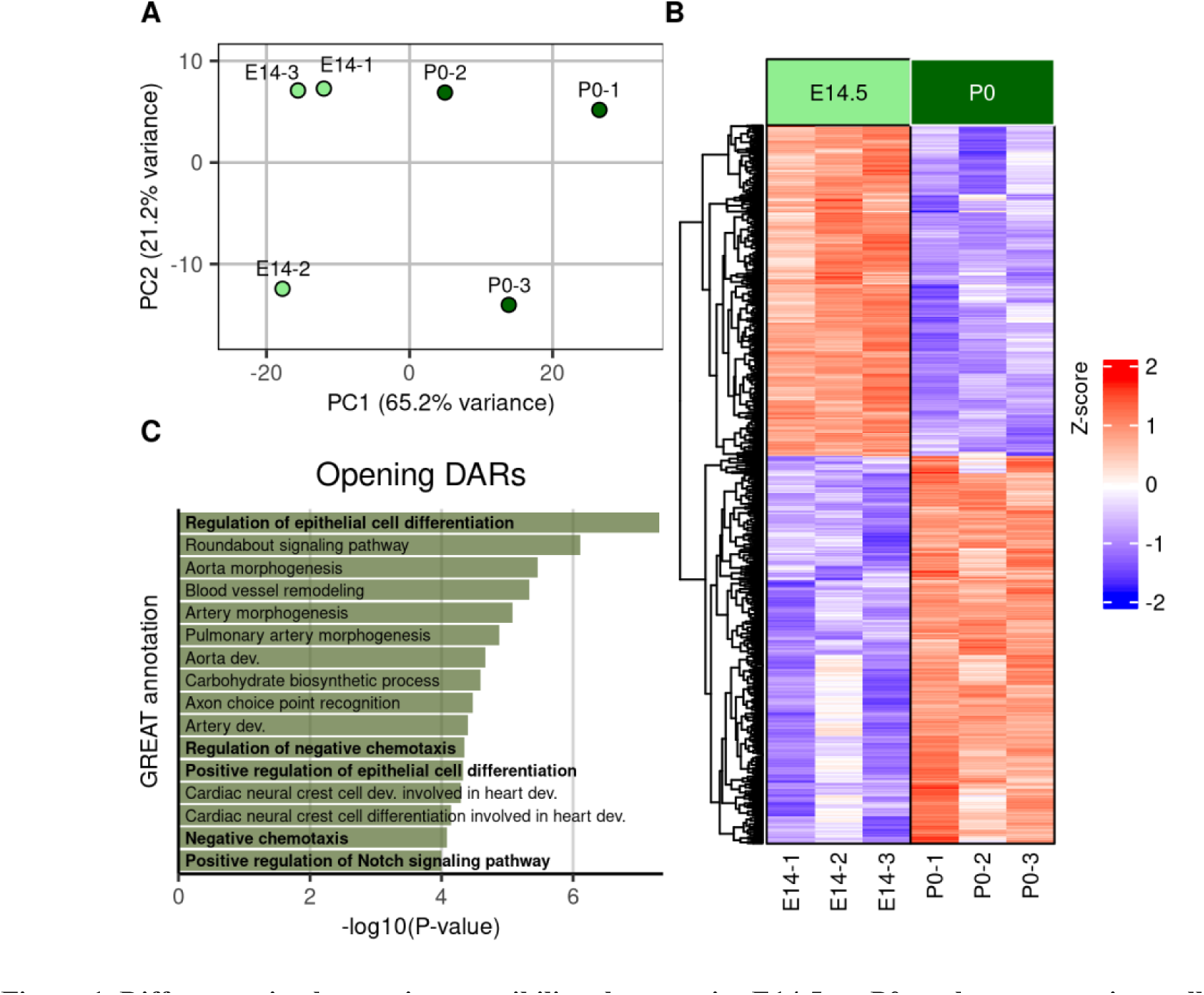
Differences in chromatin accessibility characterize E14.5 vs. P0 nephron progenitor cells. **A)** Principal component analysis of ATAC-seq samples highlights developmental time points as a major source of variation. **B)** Heatmap showing significantly changing IDR regions across ATAC-seq samples. E14.5 samples/columns are annotated in light green, and P0 samples/columns are annotated with dark green. **C)** GO Biological Process terms enriched for genomic regions that open between E14.5 and P0 according to GREAT, ranked by binomial P-value, FDR ≤ 0.001.

Transcription factor footprints were identified using HINT and matched to DNA binding proteins using motifs from the HOCOMOCO 11 database (Kulakovskiy et al., 2018). Among the 431 motifs tested, IDRs containing footprints for several transcription factors proteins associated with nephrogenesis show an increase in accessibility between E14.5 and P0. Among footprints residing in IDRs that increase in accessibility on average, the top six belong to transcription factor footprints matched to Paired protein box 8 (Pax8, 0.15), SIX homeobox 2 (Six2, 0.14), CCAAT/enhancer-binding protein δ (Cebpd, 0.13), SIX homeobox 4 (Six4, 0.13), Jun proto-oncogene (Jun, 0.12), and Fos proto-oncogene (Fos, 0.11). Footprints annotated in closing IDRs include transcription factors E2F1 and E2F4, which are known to influence cell cycle progression (Gaudet et al., 2011) and to promote cell proliferation (Wang et al., 2000) in other cellular contexts (**Supplemental figure 3**).

To evaluate the biological context of opening and closing DARs, we used the Genomic Regions Enrichment of Annotations Tool (GREAT) (McLean et al., 2010) to highlight gene sets with an enrichment of associated changing chromatin regions. With that approach we identify Gene Ontology (GO) biological processes closely tied to nephron progenitor development: in particular, the foremost enriched process among opening DARs is “Regulation of epithelial cell differentiation” (GO:0030856, Binomial Q-value = 1.5e^-5^), and “Positive regulation of epithelial cell differentiation” (GO:0030858, Binomial Q-value = 5e^-3^) is among the top 10 most significantly enriched. Both are consistent with an increase in the propensity of nephron progenitors to undergo mesenchymal-to-epithelial transitions (MET) as they differentiate (Chen et al., 2015). Other significantly enriched pathways include “Positive regulation of Notch Signaling pathway” (GO:0045747, Binomial Q-value = 2e^-5^), which is consistent with the role Notch signaling plays in promoting nephron progenitor differentiation (Chung et al., 2016).

Next, we explored chromatin accessibility in regions of the genome containing promoters and enhancers for genes with published roles in NP cell differentiation. Among these, the *Six2* transcription factor is critical for maintaining multipotency of nephron progenitor cells (Self et al., 2006), and in all samples we observed accessible chromatin surrounding its promoter and a known *Six2* enhancer ∼50kb upstream of the *Six2* transcript (O’Brien et al., 2018) **(Figure 2A, B**, blue-shaded areas). We also note regions of accessible chromatin on the promoter and a published enhancer for the protein *Gdnf* (**Figure 2D, E**), which is secreted by nephron progenitors to promote ureteric bud branching (Sanchez et al., 1996; Vega et al., 1996). This enhancer is located ∼113kb upstream of the gene promoter, and is known to be responsive to Wnt signaling in the context of developing nephron progenitors (Park et al., 2012). While the promoter for *Gdnf* does not significantly change over time, the annotated enhancer significantly increases in accessibility. We also note regions of accessible chromatin surrounding a published enhancer active in nephron progenitors for *Eyes absent 1 (Eya1)* located ∼325kb upstream of the gene’s TSS (Park et al., 2012) (**Supplemental figure 4**). Eya1 is a transcription factor which interacts with Six2 to maintain nephron progenitor multipotency (Xu et al., 2014), and its expression is essential for nephron progenitor induction (Xu et al., 1999).

**Figure 2.**
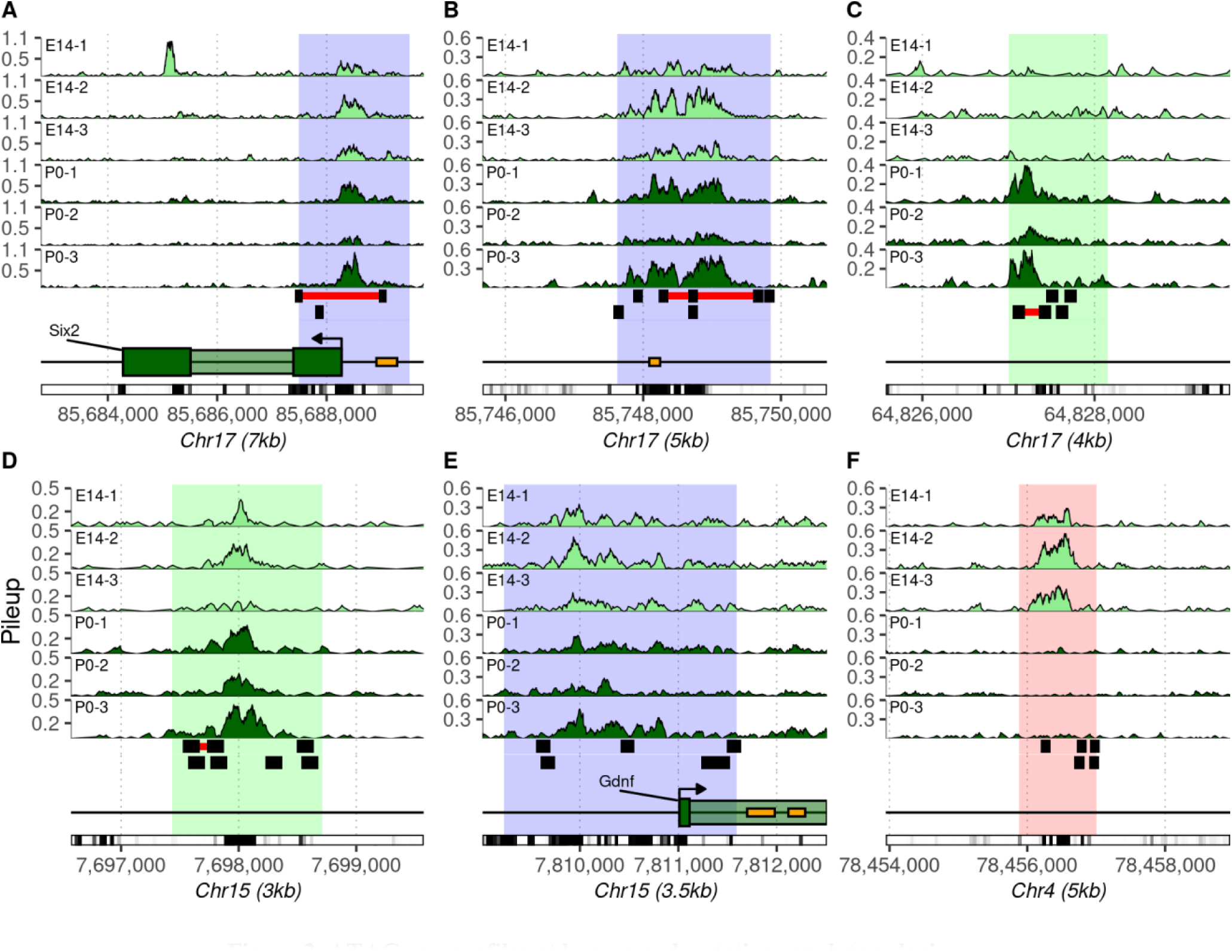
ATAC-seq profiles at known and putative regulatory loci. **A)** Accessible chromatin at the promoter of the *Six2* transcription factor. **B)** An enhancer for *Six2* with enhancer location depicted in orange(Park et al., 2012). **C)** An unannotated region of the genome on chromosome 17 exhibits a significant increase in accessibility as progenitors age. **D)** Differentially accessible enhancer for *Gdnf* with increased accessibility at P0. **E)** Promoter of *Gdnf* is accessible, but it does not significantly change over time. **F)** Unannotated intergenic DAR that is highly conserved and shows significantly reduced accessibility between E14.5 and P0. In all panels, IDR regions that are stable between time points are highlighted in blue, DARs which show increasing or decreasing accessibility with FDR controlled at 0.1 are highlighted in green and red, respectively. The top three panels in each plot show pileup in each E14.5 sample (light green), and the next three panels show pileup in each P0 sample (dark green). The remaining tracks from top to bottom indicate nucleosomal occupancy detected using NucleoATAC with nucleosomal regions annotated in black and nucleosome-depleted regions (NFRs) in red, genomic annotations, and PhastCon60 conservation. Orange blocks in annotation tracks indicate mm10 coordinates for regulatory elements identified by the FANTOM5 consortium (Andersson et al., 2014).

We observe several DARs that do not overlap known regulatory features, but encompass regions predicted to have enhancer activity for specific genes in a variety of mouse cell types and annotated in EnhancerAtlas 2.0 (Gao and Qian, 2020). In addition to the published enhancer of *Eya1* (Park et al., 2012) (**Supplemental figure 4**), a closing DAR in one of *Eya1’s* introns (**Figure 3A,B**) encompasses a region predicted to have enhancer activity on the *Eya1* gene in mouse neural progenitor and granulocyte-monocyte progenitor cells (Gao and Qian, 2020). In line with a reduction in enhancer activity, we have previously observed that expression of *Eya1* is decreased in differentiating nephron progenitors using E14.5 single-cell RNA sequencing (scRNA-seq; **Figure 3E**) (Bais et al., 2020). This potential enhancer also contains transcription factor footprints for Homeobox D10 (Hoxd10) within its peak; Hoxd10 is a transcription factor that is essential for differentiation and integration of nephron progenitors during mesenchymal-to-epithelial transitions (MET) (Yallowitz et al., 2011) and whose overexpression has been associated with Wilms tumor in humans (Royer-Pokora et al., 2010).

**Figure 3.**
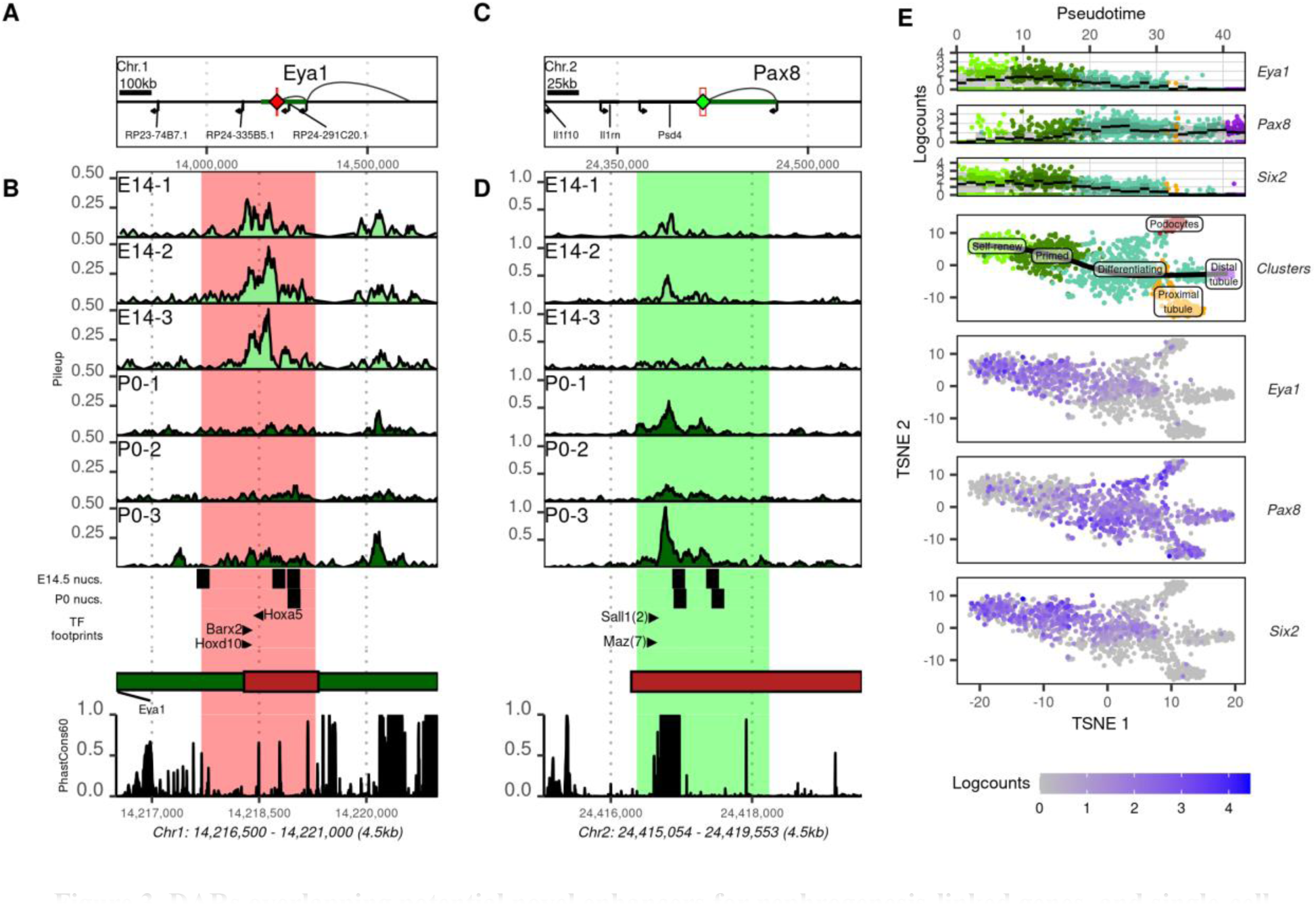
DARs overlapping potential novel enhancers for nephrogenesis-linked genes, and single-cell expression patterns of these genes among E14.5 nephron progenitors. **A)** 1MB region containing the gene body for *Eya1* (green) and a closing DAR (red diamond) that overlaps an enhancer predicted to affect *Eya1* expression. Curves originating from the *Eya1* TSS illustrate the gene’s predicted interactions with both this predicted enhancer and another published enhancer approximately 325kb upstream (**Supplemental figure 4). B)** A 4.5kb region containing the potential *Eya1* enhancer illustrated in A (red rectangle). From top to bottom, panels illustrate ATAC-seq pileup in E14.5 and P0 nephron progenitor cells (light green and dark green, respectively), E14.5 and P0 nucleosomal predictions (black), transcription factor footprints, the local gene environment, and conservation among mammals. **C)** 250kb genomic region containing the *Pax8* gene locus and a predicted enhancer downstream of the gene footprint. **D)** ATAC-seq data for a 4.5kb region containing a predicted enhancer and its overlapping DAR. **E)** Whole-kidney single-cell gene expression data for *Eya1* and *Pax8* as well as *Six2*, collected from E14.5 whole kidney samples. From top to bottom, panels 1-3 illustrate the log of gene expression in cells oriented relative to pseudotime; black bars represent mean expression, and gray boxes indicate one standard deviation. Panel 4 illustrates this pseudotime trajectory from left to right in black, plotted against a TSNE dimensional reduction of these cells and their respective cluster identities (determined using gene markers). Panels 5-7 illustrate per-cell expression levels of *Eya1, Pax8,* and *Six2* respectively.

A DAR that increases in accessibility on chromosome 24 overlaps a chromatin region with predicted enhancer activity for the transcription factor *Pax8* in cells cultured from mouse kidney, intestinal epithelia, and uterus (Gao and Qian, 2020). *Pax8* is partially redundant with *Pax2*, but at least one of these two transcription factors are required in nephron progenitors for kidney development: double mutants fail to initiate MET and eventually undergo apoptosis (Bouchard et al., 2002; Narlis et al., 2007). We previously observed that *Pax8* expression increases in differentiating and differentiated nephron progenitors (**Figure 3C, D**), and we note a significant increase in accessibility in a DAR that encompasses this regulatory feature. We observe a number of transcription factor footprints that fall within this DAR’s highly conserved accessibility peak, including two for Spalt-like transcription factor 1 (*Sall1*), a nuclear factor known to affect nephrogenesis (Kanda et al., 2014), and seven closely grouped footprints for MYC-associated zinc finger protein (*Maz*), a transcription factor which has been shown to have dosage-dependent effects on kidney development in humans and mice (Haller et al., 2018).

Other DARs identified in our ATAC-seq data may mark novel genomic regions involved in the regulatory circuitry of nephron progenitor development. Two such regions are shown in **Figure 2C** and **F**: one region on chromosome 17 and the other on chromosome 4. Both DARs fall within otherwise unannotated genomic regions, and both contain DNA sequences that are highly conserved in mammals. **Supplemental table 1** contains a complete list of the accessible regions we find as well as their respective differential accessibility statistics, annotations for genes, exons, and promoters which they overlap, and lists of key nephrogenesis-linked transcription factor footprints identified.

### Early and late nephron progenitors have distinct miRNA transcriptional profiles

Small-RNA sequencing detected 1,104 known miRNA transcripts and identified 114 miRNAs that show a significant change in expression between E14.5 and P0 samples (FDR = 0.05; **Supplemental table 2**). Half of differentially expressed miRNAs show increased expression at P0 compared with E14.5, and principal component analysis highlights greater homogeneity in miRNA expression among E14.5 samples compared with P0 (**Figure 4A**). Hierarchical clustering reveals a clear grouping of miRNAs into those of increasing and decreasing expression between E14.5 and P0, respectively **(Figure 4B**). Members of the *let-*7 family of miRNAs are among the most highly expressed miRNA detected, and we note that all family members with significant changes show increased expression at P0 (**Figure 4C**). This is in line with results showing that a reduction in *Lin28b* expression leads to a broad increase in *let-7* family expression over the course of nephrogenesis (Yermalovich et al., 2019). Other miRNAs with significantly higher expression at P0 include *miR-429-3p,* a member of the *miR-200* family known to affect podocyte differentiation (Li et al., 2016), and both *miR-125a-5p* and *miR-125b-5p.* The latter two of these are known to repress *Lin28* transcripts in different cellular environments (Chaudhuri et al., 2012; Potenza et al., 2017). miRNA with significantly lower expression at P0 include the angiogenesis- (Wang et al., 2008) and proliferation-promoting (Schober et al., 2014) *miR-126*.

**Figure 4.**
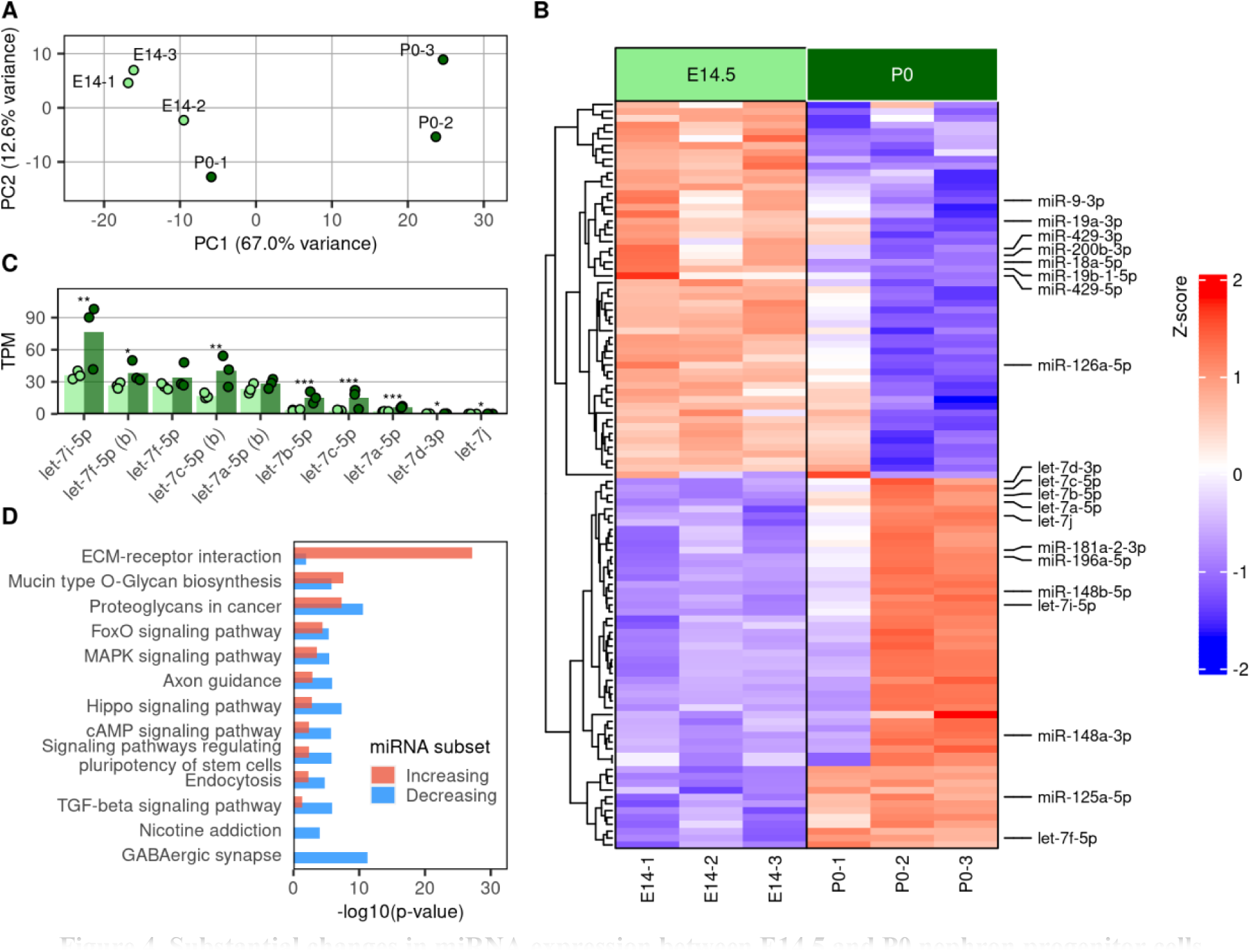
Substantial changes in miRNA expression between E14.5 and P0 nephron progenitor cells. **A)** Principal component analysis shows that developmental time point is a major contributor to miRNA expression variation. **B)** Heatmap depicting relative expression of all significantly changing miRNA. Columns containing E14.5 samples on the left are annotated in light green, and columns with P0 samples on the right are annotated in dark green. **C)** Significantly changing members of the let-7 family of miRNA are all increasing between E14.5 and P0 timepoints. **D)** Enrichment of genes from specific KEGG pathways among predicted gene targets of up- and down-regulated miRNA (red and blue bars, respectively). Minimum p-value included is 1e^-5^.

The DIANA miRPath tool enables KEGG pathway analysis of miRNA based on their respective gene targets, as predicted by microT-CDS (Vlachos et al., 2015). We find that miRNA with the greatest changes in expression between E14.5 and P0 (up or down) are those that regulate pathways with key roles in nephrogenesis (**Figure 4, Supplemental figure 5**). For instance, Tgf-β is secreted by surrounding stromal cells to promote nephron progenitor differentiation (Rowan et al., 2018), and we note reduced expression among nephron progenitor miRNA targeting the downstream effectors of the “TGF-β signaling pathway” (KEGG pathway mmu04350, p-value 1.45e^-6^). The “MAPK signaling pathway” (mmu04010) influences organization and priming of nephron progenitors for differentiation, and it regulates their interactions with the extracellular matrix (ECM) via Itgα8 (Ihermann-Hella et al., 2018). We note a significant number of up- and down-regulated miRNAs that target different genes in this MAPK signaling pathway (p-values of 3.5e^-6^ and 3.0e^-4^ for up and down lists, respectively). Finally, the KEGG pathway “Extracellular matrix (ECM) receptor interaction” (mmu04512) is the most significantly enriched pathway identified, and its constituent genes are far more significantly targeted by miRNA that increase in expression over time (p-value of 7.4e^-28^ among increasing miRNA versus 1.1e^-2^ among decreasing). ECM-associated genes are targeted by some of the most highly expressed and significantly increasing miRNA we detect, including *miR-196a-5p*, *miR-148a-3p, miR-125a-*5p, and *let-7f-5p*. In particular, *miR-196a-5p* is the highest expressed miRNA in our samples, and it sees an 8-fold increase between E14.5 and P0. Gene transcripts that were the predicted targets of miRNA (using TargetScan (Agarwal et al., 2015)) with increased expression at P0 include *Bach2*, a proposed molecular link between the MAPK/AP1 and Six2/β-catenin pathways for self-renewal and differentiation in nephron progenitors, respectively (Hilliard et al., 2019).

### Topologically associated domains help identify potential enhancer—miRNA relations

Topologically associated domains (TADs) are megabase-scale regions of higher-order chromatin organization that result from the packaging of chromatin into the nucleus in three-dimensional space (Dixon et al., 2012), and enhancer-promoter interactions are typically restricted to elements within the same TAD (Symmons et al., 2014). We approximated TADs from nephron progenitors using annotations generated from mouse embryonic stem cells (ESCs) (Wang et al., 2018). To screen for possible cis-regulatory regions of miRNA, we correlated changes in chromatin accessibility with differentially expressed miRNA within the same TAD. Overall, 42,778 unique accessible regions (about 92%) are located inside of annotated TADs, and of these 30,626 (about 72%) do not overlap a known promoter. This includes 2,103 DARs, with 1,180 and 923 opening and closing over time, respectively. Fifteen opening DARs share a TAD with one or more miRNAs with increased expression between E14.5 and P0, and nine closing DARs share a TAD with one or more miRNA with decreased expression. **Figure 5** summarizes this approach and illustrates distances between DARs and their matched miRNA, which range from 27 kb to over 1 mb (average: 447 kb). **Table 1** details sets of possible DAR – miRNA relationships, with one DAR listed per matched miRNA.

**Table 1.** miRNA exression changes, matched to one or more DAR in the same TAD.

| **miRNA name** | **log2FC** | **P-adjusted** | **Closest DAR** | **Distance (kb)** |
| --- | --- | --- | --- | --- |
| *let-7d-3p* | 1.6 | 4.2e-2 | Chr13:49,716,724-49,717,659 | 1,181 |
| *let-7c-5p* | 2.6 | 5.6e-4 | Chr16:77,719,058-77,720,407 | 119 |
| *let-7a-5p* | 2.6 | 3.4e-4 | Chr13:49,716,724-49,717,659 | 1,178 |
| *miR-9-3p* | -3.5 | 3.8e-3 | Chr13:83,987,514-83,989,301 | 249 |
| *miR-125b-5p* | 1.7 | 3.0e-2 | Chr16:77,719,058-77,720,407 | 73 |
| *miR-137-3p* | -6.1 | 3.0e-3 | Chr3:118,674,424-118,676,014 | 240 |
| *miR-181a-2-3p* | 1.8 | 4.1e-2 | Chr2:38,722,003-38,722,860 | 130 |
| *miR-182-5p* | 1.7 | 3.0e-2 | Chr6:30,359,230-30,360,992 | 193 |
| *miR-183-5p* | 2.0 | 3.0e-2 | Chr6:30,359,230-30,360,992 | 189 |
| *miR-186-3p* | -2.5 | 3.0e-2 | Chr3:157,078,826-157,081,050 | 463 |
| *miR-455-3p* | 2.1 | 1.3e-2 | Chr4:63,224,743-63,226,213 | 31 |
| *miR-467a-3p* | -2.4 | 1.9e-2 | Chr2:10,614,913-10,616,335 | 131 |
| *miR-467a-3p (b)* | -3.0 | 6.0e-4 | Chr2:10,614,913-10,616,335 | 119 |
| *miR-486b-5p* | 2.2 | 2.6e-2 | Chr8:23,037,345-23,039,410 | 103 |
| *miR-486a-5p* | 2.2 | 2.4e-2 | Chr8:23,037,345-23,039,410 | 103 |
| *miR-486a-3p* | 5.0 | 4.2e-2 | Chr8:23,037,345-23,039,410 | 103 |
| *miR-676-3p* | 2.6 | 1.6e-3 | ChrX:99,858,825-99,859,557 | 522 |
| *miR-879-3p* | -4.8 | 2.1e-2 | Chr5:9,485,240-9,487,070 | 109 |
| *miR-1298-5p* | -6.5 | 2.4e-3 | ChrX:144,407,791-144,408,724 | 2,656 |
| *miR-1843b-3p* | 2.7 | 2.9e-2 | Chr1:160,083,545-160,084,967 | 743 |
| *miR-3970* | 2.2 | 4.7e-2 | Chr19:33,129,533-33,131,028 | 26 |
| *miR-5099* | 1.6 | 4.2e-2 | Chr12:36,725,420-36,726,957 | 89 |
| *miR-5108* | 7.3 | 2.4e-2 | Chr10:61,964,959-61,965,989 | 190 |

**Table 2.**
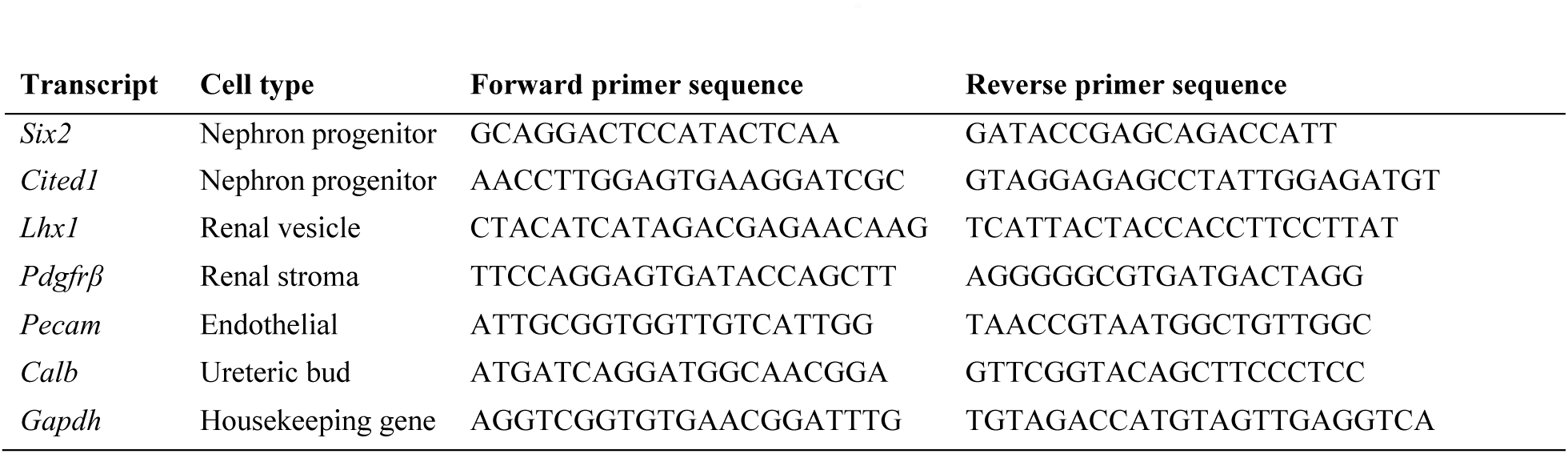
RT-PCR primers.

**Figure 5.**
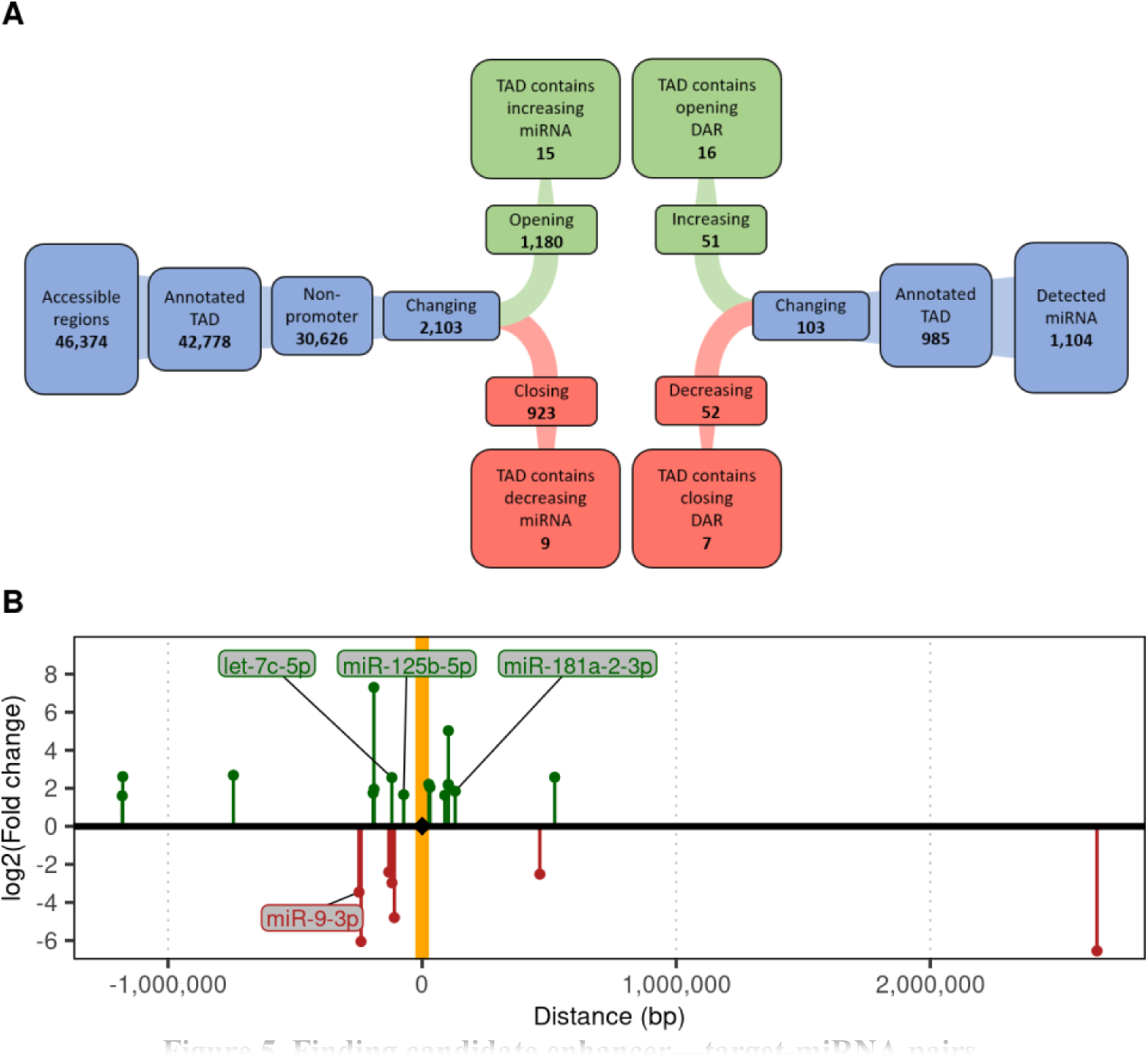
Finding candidate enhancer—target-miRNA pairs. **A)** Flow chart denoting how regions of accessibility were prioritized for possible regulation of miRNA expression. **B)** Genomic locations of each screened miRNA relative to their candidate enhancer (orange vertical stripe). The scale of their log2 fold change is shown on the y-axis, and miRNA with increasing or decreasing expression are highlighted in green and red, respectively.

Among the miRNA-enhancer pairs annotated, we note three of particular interest. First, an opening DAR has been identified 120 kb downstream of *let-7c-5p* and 73 kb downstream of *miR-125b-5p* (**Figure 6A**)*. Let-7c-5p* is one of the highest-expressed *let-7* miRNA we detect, and the *let-7* family is known to also repress Lin28b (Slack and Ruvkun, 1997; Zhu et al., 2011). *miR-125b-5p* has also been shown to repress *Lin28* transcripts in mouse hematopoietic and progenitor cell types (Chaudhuri et al., 2012). Within this putative enhancer, we detect DNA binding footprints for transcription factors known to play important roles in kidney development, including *Sox9*, a transcription factor known to mark highly proliferative “progenitor like” cells during kidney repair (Kang et al., 2016; Kumar et al., 2015). Second, *miR-9-3p* is the reverse complement to *miR-9-5p*, a miRNA which has recently been shown to protect from kidney fibrosis by targeting metabolic pathways (Fierro-Fernández et al., 2020). We detect a greater than 10-fold decrease in *miR-9-3p* transcripts between E14.5 and P0 nephron progenitors, and we have identified a potential regulatory element approximately 249kb away **(Figure 6B**). Finally, we matched *miR-181a-2-3p* to a possible intergenic cis-regulatory locus 131kb upstream on chromosome 2, where we note a transcription factor footprint for the transcription factor *Wt1*, a key regulator of nephron progenitor survival (Kreidberg et al., 1993) (**Figure 6C**).

**Figure 6.**
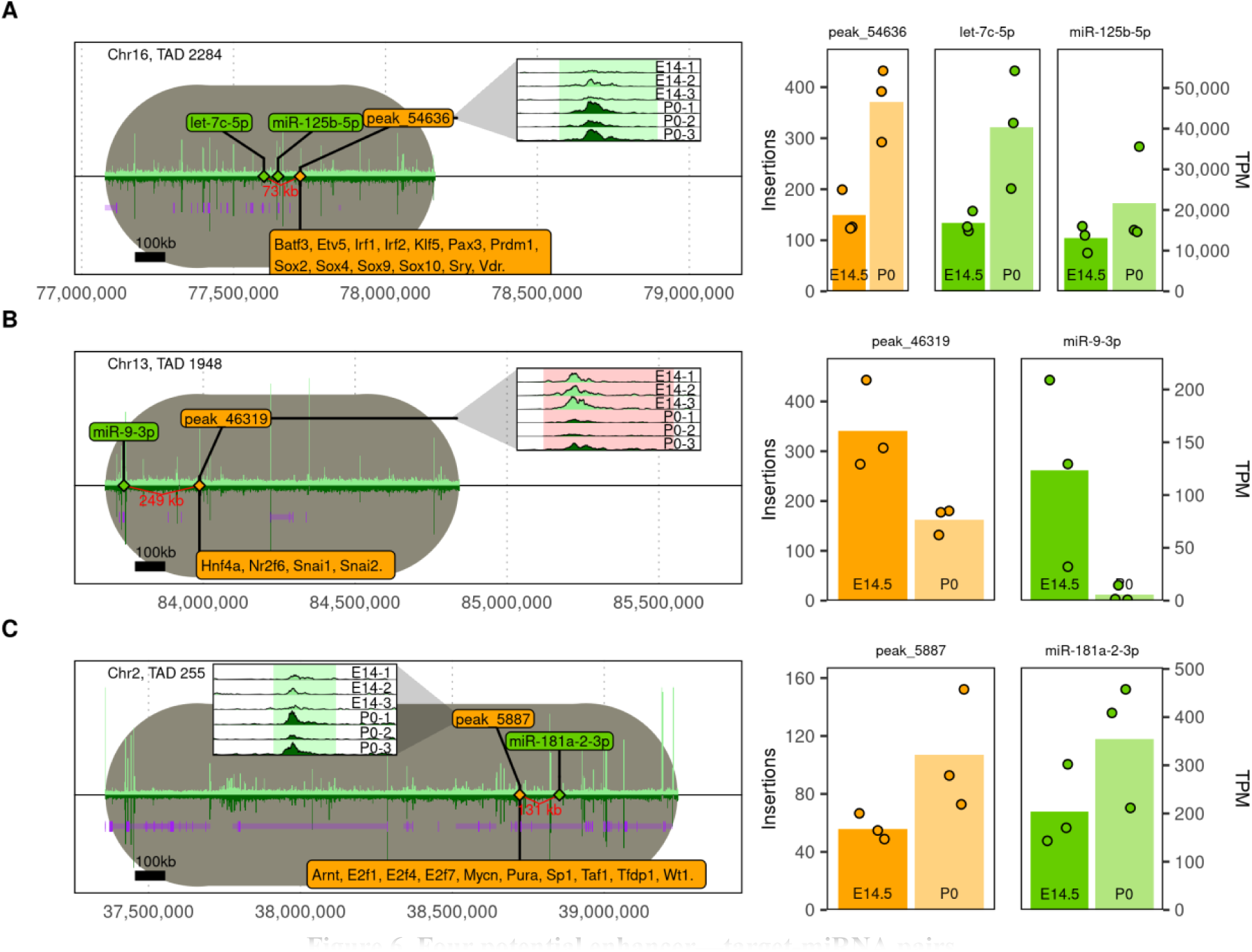
Four potential enhancer—target-miRNA pairs. TADs are highlighted in gray, aligned by starting points and drawn in the same scale to illustrate relative size. Known genes from Ensembl are shown in purple, and darker regions indicate exons. Relative accessibility at E14.5 and P0 are shown as bar plots above and below the horizontal black line, respectively, as well as in inset plots illustrating pileup in the specified DAR. Transcription factor footprints identified by HINT are listed in orange banners. Bar plots in separate panels on the right indicate miRNA expression (green) and normalized Tn5 insertion events in proposed enhancer regions (orange). **A)** The let-7 family member *let-7c-5p* is very highly expressed in nephron progenitor cells and is matched to “peak_54636” 119kb away in an intergenic region. **B)** *miR-9-3p* is matched to a DAR 249kb away. **C)** miR-181a-2-3p is located 131kb away from “peak_5887,” a possible intergenic enhancer.

Combined, these data demonstrate that changes intrinsic to nephron progenitors and dependent on the stage of nephrogenesis are reflected in the progenitor cell’s chromatin environment and in its miRNA transcriptome. These data can be utilized to identify regulatory features or miRNA species of interest in the nephron progenitor cell, as well as the gene transcripts they may ultimately affect.

## Discussion

Our data demonstrate that E14.5 and P0 nephron progenitor cells are characterized by distinct chromatin environments and miRNA transcriptomes. We find that many of the differences in each (DNA accessibility and miRNA expression) implicate parallel developmental processes, including cell migration and epithelial cell development. We discussed several concrete examples where our data implicate novel gene-regulatory loci, and we pointed out 33 potential miRNA-enhancer interactions, including for the miRNAs *let-7-5p*, *miR-125b-5p*, *miR-181a-2-3p*, and *miR-9-3p*. We expect our data to be useful for further exploration and in the context of future functional genomics studies in the context of nephron progenitor cells.

We describe two putative novel enhancers for *Eya1* and *Pax8*, based on *(i)* their proximity to these gene promoters, *(ii)* their respective changes in chromatin accessibility during development being in-line with pseudotime gene expression changes during differentiation, *(iii)* footprints of kidney development transcription factors, *(iv)* sequence conservation, and *(v)* evidence of enhancer function in other tissues / cell types from Enhancer Atlas2.0 (Gao and Qian, 2020). We observe a decrease in chromatin accessibility for the *Eya1* enhancer, which is consistent with *Eya1*’s role in promoting nephron progenitor multipotency, and with reports that loss of *Eya1* results in the early epithelialization of the progenitor population (Xu et al., 2014, 1999). *Pax8* and its homolog *Pax2* contribute to lineage specification of nephron progenitors (Bouchard et al., 2002), and nephron progenitors without at least one of these genes will not undergo MET (Narlis et al., 2007). The increased chromatin accessibility we detect for the putative *Pax8* enhancer is consistent with increased *Pax8* expression in renal epithelia as nephrons differentiate.

Enrichment analysis (McLean et al., 2010) of DARs revealed “Regulation of Epithelial Differentiation” as the most significantly enriched term among opening chromatin regions, indicating that the DARs we detect may contain regulatory features driving or responding to signals for differentiation and development of nephron progenitor cells into epithelial cell types. Among the genes nearby such DARs are *Yes-associated protein 1* (*Yap1*), a transcriptional coactivator in nephron progenitors that is required for the induction of nephron differentiation (Reginensi et al., 2013), and *Hairy/Enhancer of Split* (*Hes1*), a transcriptional repressor and downstream effector of Notch signaling whose expression affects the patterning of epithelial cells during morphogenesis (Piscione et al., 2004). Notch signaling is also among the most significantly enriched GO terms identified by opening DARs, and in nephron progenitors this pathway is known to repress *Six2* expression and promote nephron progenitor cell differentiation (Chung et al., 2016). Among this pathway’s genes implicated by nearby opening chromatin regions is *Jagged1* (*Jag1*), a key ligand in Notch signaling whose expression has been reported to be inversely correlated with expression of *Six2* and *Hes1* (Chung et al., 2016). Finally, we observe enrichment for genes associated with blood vessel remodeling and morphogenesis in opening DARs, although this appears to be a consequence of genes shared by these developmental pathways with epithelial cell differentiation, including *Jag1*, *Eya1, Hes1, Forkhead box C1* (*Foxc1*), and *Hairy/enhancer of split related with YRPW motif 2* (*Hey2*).

Changes in the nephron progenitor’s chromatin environment coincide with changes in the miRNA transcriptome. We observe broad increases in expression of the *let-7* family as progenitors age, which is consistent with previous reports: Lin28b and the *let-7* family of miRNAs are mutually antagonistic and form a bistable switch that governs *let-7* expression in mammals (Zhu et al., 2011), and in nephron progenitors the switch toward increased *let-7* expression directly affects the timing of the cessation of nephrogenesis (Yermalovich et al., 2019). We also observe increased expression between E14.5 and P0 in for *miR-125b-5p,* which is also known to repress expression of *Lin28b* in other cell contexts (Chaudhuri et al., 2012; Potenza et al., 2017). These two miRNAs are among 114 others that are differentially expressed between E14.5 and P0 in nephron progenitors, which suggests that the role of miRNAs in nephron progenitor development is not limited to *let-7*/*Lin28b*. For instance, miRNAs which decrease in expression include two members of the *miR-200* family, which has been reported to play a role in podocyte differentiation (Li et al., 2016): *miR-200b-3p* and *miR-429-3p.* Further on, we observe a decrease in the expression of *miR-126a-5p*, which is known to promote vasculogenesis (Schober et al., 2014; Wang et al., 2008). This may reflect the common mesenchymal lineage of vasculogenic and nephron progenitors, and commitment of a subset of these cells to become nephron progenitors.

Pathway analyses of predicted gene targets of changing miRNAs using DIANA miRPath (Vlachos et al., 2015) show enrichment for a variety of known KEGG pathways, including MAPK and TGF-β signaling, both key regulators of nephron progenitors (Hilliard et al., 2019; Ihermann-Hella et al., 2018). Several other KEGG pathways are also overrepresented among the predicted gene targets of up-regulated miRNAs, including pathways related to extracellular matrix, actin cytoskeleton, and focal adhesion: all crucial facets of cellular migration, differentiation, branching, attachment, polarization, and proliferation (Rozario and Desimone, 2010). ECM-associated genes are targeted by some of the most highly expressed and significantly increasing miRNAs we detect, including the highest expressed transcript we detect, m*iR-196a-5p. miR-196a-5p* is known to target *Col1a1, Col1a2,* and *Col3a1*—genes which are up-regulated in *Pax2*-deficient nephron progenitors that undergo transdifferentiation into the renal stroma (Naiman et al., 2017). Thus, increased *miR-196a-5p* expression may have a role in bolstering lineage boundaries as progenitors age.

By combining observed changes in miRNA expression with changes in the chromatin environment, we identify several novel miRNA—enhancer pairs. The miRNAs *miR-125a-5p* and *let-7c-5p*, for instance, see a significant increase in expression between E14.5 and P0, and this increase in expression is recapitulated by an increase in accessibility of a DAR (chr16:77,719,058 77,720,407) in the same TAD. Within this DAR we observe several transcription factor footprints for members of the *Sox* family, which are known regulators of cell fate in stem and progenitor cells (Sarkar and Hochedlinger, 2013). *Sox9* in particular marks highly proliferative multi-lineage progenitor cells in mouse kidneys (Kang et al., 2016; Kumar et al., 2015). This DAR may denote an enhancer that affects expression of *miR-125a-5p* and/or *let-7c-5p,* which in turn could repress *Lin28*, thereby promoting differentiation. We observe a significant increase in P0 of the expression of *miR-181a*, which is known to modulate renin expression in humans (Marques et al., 2015) and has been considered a therapeutic target in humans to reduce nephrotoxicity due its ability to repress BIRC6 and apoptosis (Liu et al., 2018). This miRNA is matched to a possible enhancer region 131kb away which becomes significantly more accessible over the same time frame. Finally, *miR-9-3p* is located 249kb from a closing DAR, and we observe a ten-fold decrease in expression between E14.5 and P0. *miR-9-3p*’s reverse complement, *miR-9-5p*, is known to protect from kidney fibrosis (Fierro-Fernández et al., 2020), and we observe transcription factor footprints for the *Wnt* signaling-responsive proteins Snail (Snai1) and Slug (Snai2) in the associated DAR. Snai1 and Snai2 have been shown to promote carcinoma metastasis in mouse skin cancer cells (Olmeda et al., 2008) and to promote epithelial-to-mesenchymal transitions (EMT) in cancer patients (Sundararajan et al., 2019), while *miR-9* is known to promote angiogenesis and migration of tumor endothelial cells (Zhuang et al., 2012). A mechanism may exist in which increased Wnt signaling results in reduced expression of *Snai1* and *Snai2*, which in turn reduces *miR-9-3p* expression to support nephron progenitor differentiation.

Our study comprehensively identified changes in the chromatin landscape and the miRNA transcriptome of nephron progenitors during embryonic development, and we have identified candidate cis-regulatory regions and micro RNA that likely play a role in the cessation of nephrogenesis. Enrichment analyses for both the ATAC-seq and smRNA-seq data provide enticing clues of the mechanisms that may be involved, indicating modulating pathways that regulate nephron progenitor differentiation, such as Notch and TGF-β. We also report two novel putative regulatory elements for *Eya1* and *Pax8*, genes known to be important in kidney development, and potential regulatory sequences for key miRNA that are implicated and known in the cessation of nephrogenesis.

## Methods

### Mouse strains

Wildtype CD-1 time-mated pregnant females were obtained from Charles River Laboratories, Inc. (Wilmington, MA, USA, RRID:MGI:5659424) and kidneys were collected from litters at embryonic day E14.5 or P0. All animals were housed in the vivarium at the Rangos Research Center at the UPMC Children’s Hospital of Pittsburgh (Pittsburgh, PA, USA) and all animal experiments were carried out in accordance with the policies of the Institutional Animal Care and Use Committee at the University of Pittsburgh. The sex for each embryo or pup was identified by performing PCR on genomic DNA isolated from tail clippings using the following primers for the Y-chromosome gene *Sry*: SryF 5ʹ-GATGATTTGAGTGGAAATGTGAGGTA-3ʹ and SryR 5ʹ-CTTATGTTTATAGGCATGCACCATGTA-3’, as previously published (McFarlane et al., 2013).

### Nephron progenitor isolation

Kidneys were dissected from litters of 8-12 E14.5 embryos or P0 pups, and each litter was considered one sample. One kidney per sample was used for total RNA isolation to be used as a “whole kidney control” in subsequent analyses (see below). A total of three samples were collected at each time point. Cortical cells were then isolated from each sample using magnetic-activated cell sorting (MACS) as previously published (Brown et al., 2011). Briefly, kidneys underwent partial digestion with 2mM collagenase A (Roche 11088793001) and 3.5mM pancreatin (Sigma P1625) in phosphate-buffered saline (PBS) for 15 min at 37°C. The digestion reaction was halted using cold 100% fetal bovine serum (FBS), and the cell suspension was collected and resuspended in cold Dulbecco’s phosphate-buffered saline (DPBS) with 1mM phenylmethylsulfonyl fluoride (PMSF) protease inhibitor (Sigma-Aldrich 10837091001). The cells were washed and resuspended in ice-cold isolation buffer consisting of 2% FBS in PBS, then incubated with magnetic beads (Dynabeads FlowComp Flexi kit, ThermoFisher 11061D) biotinylated to antibodies for Integrin alpha 8 (Itgα8, R&D Systems AF4076) using the DSB-X Biotin Protein Labeling Kit (ThermoFisher D-20655). Nephron progenitors bound to these beads were isolated as previously published (Hemker et al., 2020) and pooled across kidneys for each sample. An aliquot of 50,000 nephron progenitor cells was immediately processed through the chromatin accessibility library preparation protocol (see below), and total RNA was extracted from the remaining cell suspension using the Qiagen miRNeasy Mini Kit (Qiagen 217004).

To confirm enrichment of nephron progenitor cells relative to other cell types, total RNA was isolated from nephron progenitors as well as from whole kidneys (whole kidney control, see above), and quantitative reverse-transcription PCR (qRT-PCR) for the following markers was performed: nephron progenitor markers *Six2* and *Cited1* (Huang et al., 2016), and for *Lhx1, Pdgfrβ, Pecam* and *Calb* that mark renal vesicle (Brunskill et al., 2014), renal stroma (Chen et al., 2015), endothelial (Kobayashi et al., 2008), and ureteric bud (Cebrian et al., 2014b) cells, respectively; also see **Supplemental figure 1**. *Gapdh* (Barber et al., 2005) was used as housekeeping gene and relative quantification was calculated via the 2^-ΔΔCt^ method (Livak and Schmittgen, 2001).

### Chromatin accessibility library preparation

Approximately 50,000 nephron progenitor cells were used to generate each sequencing library for the assay for transposase-accessible chromatin (ATAC-seq) using the Nextera DNA Flex Library Prep Kit (Illumina FC-121-1030) with modifications according to a previously published method (Buenrostro et al., 2013). In brief, cells were suspended in an ice-cold non-ionic lysis buffer consisting of 10mM Tris, 10mM NaCl, 3mM MgCl2, 1mM PMSF protease inhibitor, and 0.05% Triton X-100 by volume for 10 minutes to lyse cell membranes while leaving nuclear membranes intact. Intact nephron progenitor nuclei were resuspended in transposition buffer containing the transposase enzyme and incubated at 37°C for 30 minutes. Free genomic DNA released by the transposition process was purified using the MinElute PCR Purification kit (Qiagen 28004) and then indexed using forward (i7) and reverse (i5) index primers from the Nextera Index Kit (Illumina FC-121-1011). Index ligation and fragment amplification were achieved using the method’s PCR amplification thermal cycling program.

To determine the optimal number of amplification cycles required for the ATAC-seq library, a qPCR side reaction was performed using SsoAdvanced SYBR Supermix (BioRad 1725274) and a 96-well C100 Thermal Cycler (Bio-Rad) to calculate the normalized reporter value (Rn) for each cycle. The cycle number at which the reaction reached one-third of its maximum fluorescence was identified as the optimal number of amplification cycles remaining, and the remaining volume of the indexed ATAC-seq transposition reaction was subjected to that number of additional cycles in the final amplification program. Final libraries were purified using Ampure XP magnetic beads (Beckman Coulter A63881).

### Chromatin accessibility sequencing

Paired-end sequencing of the ATAC-seq library was performed on an Illumina NextSeq500 by the Health Sciences Sequencing Core at UPMC Children’s Hospital of Pittsburgh, multiplexed with library concentrations expected to yield 90 million paired end reads per sample. Reads were quality trimmed using TrimGalore (BabrahamLab, 2014) (version 0.4.3) in “--paired” mode and with otherwise default settings. Reads were aligned to the mm10 genome (Church et al., 2011) using Bowtie2 (Langmead et al., 2009) (version 2.3.1) with the settings “--local -q -X 2000 --m”. Reads resulting from PCR duplicates of the same DNA fragment were marked using Picard Tools’ (Broad Institute, 2016) MarkDuplicates function (version 2.10.9) with default settings. Reads that mapped ambiguously to more than one location in the genome were randomly assigned to one of these locations if there were fewer than four possibilities, and were otherwise eliminated (ENCODE, n.d.) (using a Python script available at https://github.com/kundajelab/atac_dnase_pipelines, (Lee, 2017) with the settings “--paired-end -k”). Samtools (Li et al., 2009) (version 1.3.1) was used for filtering out duplicated, unmapped, orphaned, and mitochondrial reads, as well as the sorting and indexing of BAM files. Quality control of ATAC-seq data followed the ENCODE project’s ATAC-seq guidelines (ENCODE, n.d.), and sample libraries were randomly down-sampled to each contain 31 million paired-end reads in order to normalize for differences in library sizes.

### Identifying accessible chromatin regions

BAM files containing filtered ATAC-seq reads were converted into paired-end BED format using Bedtools’(Quinlan and Hall, 2010) (version 2.26.0) ‘bamtobed’ function with settings “-bedpe - mate1”. Regions of accessible chromatin were identified using the Model-based Analysis of ChIP-seq tool (MACS2, version 2.1.1.20160309) broad peak calling algorithm (Zhang et al., 2008), with settings “--format BEDPE --g mm -p 0.01 --broad --shift 37 --extsize 75 --keep-dup all --nomodel". To identify high-confidence accessible regions, pair-wise comparisons (of each combination of replicates from a given time point) were performed and the irreproducible discovery rate (IDR) (Li et al., 2011) was calculated using a publicly available Python implementation (GitHub https://github.com/nboley/idr) (Boleu et al., 2015) with settings “--input-file-type broadPeak -- rank p.value --soft-idr-threshold 0.1 --output-file-type bed”. Specifically, for each time point a concatenated BED file of all accessible regions was submitted through the “--peak-list” argument for each comparison (for instance, when comparing two E14.5 samples, all E14.5 broadPeaks between all three replicates are used). Regions of accessibility that were consistent by IDR in at least two pair-wise replicate comparisons (with a minimum of 20% overlap) were combined into a unified set of high-confidence accessible regions (“IDR regions”).

### Changes in accessible chromatin regions

The accessibility of each IDR region within a given sample was quantified by counting the number of transposition events that occur while normalizing for sequence GC-content across samples. This was achieved using a custom script (available at the accompanying software repository, see supplemental data). Changes in this accessibility were then determined by comparing these values across time points using the Limma-voom (version 3.42.2) software package (Ritchie et al., 2006). To account for embryo sex, the fraction of female embryos in each sample was included as a cofactor in linear models used for differential accessibility analysis, and it was removed as a batch effect in visualizations using the Limma package’s removeBatchEffect function. Differentially accessible regions of chromatin (DARs) were deemed to be significantly opening (increased accessibility between E14.5 and P0) or closing (decreased accessibility between E14.5 and P0) when controlling the false discovery rate at 10%. Functional enrichment analyses of opening and closing genomic regions was determined by submitting respective coordinates to the Genomic Regions Enrichment of Annotations Tool (GREAT, available at http://great.stanford.edu/public/html/) (McLean et al., 2010). Annotations of known and predicted enhancers were retrieved from the FANTOM5 (Andersson et al., 2014), VISTA (Visel et al., 2007), and EnhancerAtlas 2.0 (Gao and Qian, 2020) repositories.

### Transcription factor footprints

Transcription factor footprints were identified by pooling all BAM files from the same time point, then running the Regulatory Genomics Toolbox’s Hmm-based Identification of TF Footprints (HINT) software (version 0.12.3), specifically the footprinting model for ATAC-seq data (available at www.regulatory-genomics.org) (Li et al., 2018). This software was run with settings “rgt-hint footprinting --organism=mm10 --atac-seq --paired-end”. Footprints were then annotated with known mouse binding motifs from the HOCOMOCO 11 database (Kulakovskiy et al., 2018) using HINT’s motif matching function with the settings “rgt-motifanalysis matching -- organism=mm10”. Footprints with HINT scores below 10 were excluded. Finally, differential activity patterns among motif-matched footprints was measured using the RGT-HINT program’s differential function, with settings “rgt-hint differential --organism=mm10 --bc --nc 12 --output- profiles”. Significantly changing activity levels were identified based on a p-value cutoff of 0.05 (Friedman-Nemeny method).

### Nucleosomal configuration

Aligned reads from the same time point were pooled into a single BAM file, and the NucleoATAC package (version 0.3.4) was used to identify patterns in read lengths suggestive of both bound nucleosomal dyads and of nucleosome-free regions (NFRs) (Schep et al., 2015). This software was executed using default settings. Nucleosomes were annotated as the 146bp region centered around each called nucleosome dyad.

### Single-cell RNA expression

Single-cell gene expression data was processed for a previous manuscript (BioRxiv https://www.biorxiv.org/content/10.1101/2020.09.16.300293v1) (Bais et al., 2020), and was retrieved from the Gene Expression Omnibus (GEO ID GSE158166). Processed data incorporated into (**Figure 3)** include cell types, gene expression levels, low-dimensional (tSNE) embeddings, and pseudotime annotations relative to the trajectory established between clusters of proliferating and differentiating nephron progenitors. Data were accessed and organized using the SingleCellExperiment R package (Lun and Risso, 2019).

### Small RNA sequencing and analysis

Total RNA isolated from the nephron progenitor cells remaining after removing the ATAC-seq cell fraction was submitted to the Health Sciences Sequencing Core at UPMC Children’s Hospital of Pittsburgh for library preparation using the QIAseq miRNA library preparation kit (Qiagen 331502). Single-end sequencing of smRNA-seq libraries was performed on an Illumina NextSeq500, and libraries were sequenced to a depth of approximately 50 million single-ended reads per library. Sequenced reads were then aligned to the mouse mm10 genome using the Rsubread package (Liao et al., 2019) (version 2.0.1), and annotated to known miRNA listed in miRBase (version 22) (Kozomara and Griffiths-Jones, 2014) with the Rsubread package’s featureCounts function. Differential expression of known miRNA was measured using the DESeq2 R package (version 1.26.0) (Love et al., 2014). The fraction of aligned reads annotated to known miRNA as well as the fraction of female embryos per sample were included as cofactors in the testing model submitted to DESeq2. Differentially expressed miRNA were then identified while controlling the false discovery rate at 5%, and their direction of change (increasing vs. decreasing) was determined based on their fold change value (“increasing” indicating higher expression at P0 compared with E14.5). Enrichment of regulatory pathways among predicted gene targets of miRNA with significantly increasing and decreasing expression were calculated using DIANA tools miRPath version 3.0 (Vlachos et al., 2015). Predicted target transcripts for miRNA were retrieved from the TargetScan (Agarwal et al., 2015) and miRDB (Chen and Wang, 2020) repositories.

### Screening for regulatory elements affecting miRNA expression

To screen for potential enhancers of miRNA, we annotated miRNA and DARs with the topologically associated domains (TADs) in which they are located based on mm10 annotations from mouse embryonic stem cells, downloaded from the 3D Genome Browser (available at http://3dgenome.fsm.northwestern.edu/) (Wang et al., 2018). Regions of chromatin accessibility and miRNA were considered candidate enhancer—miRNA pairs if they occupied the same TAD and showed consistent changes in accessibility and expression (increasing accessibility with increasing miRNA expression and vice versa). DARs were only considered as possible “regulatory features” if they did not overlap a known promoter or exon end. Potential pairs of miRNA and putative regulatory elements were prioritized if they 1) fell within the same TAD and 2) experienced significant changes between E14.5 and P0 (increasing miRNA expression and increasing IDR region accessibility, and vice versa).

## Acknowledgements

Shelby Hemker and Abha Bais contributed to experimental design. Sequencing for ATAC-seq and smRNA-seq was performed by the Health Sciences Sequencing Core at UPMC Children’s Hospital of Pittsburgh

## Data availability

Predicted mouse enhancers were retrieved from FANTOM5 (DOI: 10.1038/nature12787; file available [here](https://fantom.gsc.riken.jp/5/datafiles/latest/extra/Enhancers/mouse_permissive_enhancers_phase_1_and_2.bed.gz)), Vista (DOI: 10.1093/nar/gkl822; file available [here](https://enhancer.lbl.gov/cgi-bin/imagedb3.pl?page_size=100;show=1;search.result=yes;page=1;form=search;search.form=no;action=search;search.sequence=1)), and EnhancerAtlas (DOI: 10.1093/nar/gkz980; mouse files available [here](http://www.enhanceratlas.org/downloadv2.php)) databases. The mouse mm10 genome and its annotations were downloaded via the Illumina iGenomes project (available [here](http://igenomes.illumina.com.s3-website-us-east-1.amazonaws.com/Mus_musculus/UCSC/mm10/Mus_musculus_UCSC_mm10.tar.gz)). The overchain file for lifting mm9 coordinates to mm10 was retrieved from the UCSC genome browser (DOI: 10.1101/gr.229102; file available [here](http://hgdownload.soe.ucsc.edu/goldenPath/mm9/liftOver/mm9ToMm10.over.chain.gz)). TAD annotations from mouse embryonic stem cells were retrieved from the 3D Genome Browser (DOI: 10.1186/s13059-018-1519-9; file available [here](http://3dgenome.fsm.northwestern.edu/downloads/mm9.TADs.zip)). Predicted miRNA transcript targets were retrieved from TargetScan (DOI: 10.7554/eLife.05005, file available [here](http://www.targetscan.org/mmu_72/mmu_72_data_download/Conserved_Family_Conserved_Targets_Info.txt.zip)), and from miRDB version 6 (DOI: 10.1093/nar/gkz757, file available [here](http://mirdb.org/download/miRDB_v6.0_prediction_result.txt.gz)). ATAC-seq data including IDR/DAR locations, annotations, and differential enrichment results are available on GEO (GSE168339), as are smRNA-seq data including annotations and differential expression results (GSE168342). Computer code used to process and analyze data, and to generate figures is available at https://bitbucket.org/clugstonA/mirna_enhancers.

## Supplemental figures

**Supplemental figure 1.**
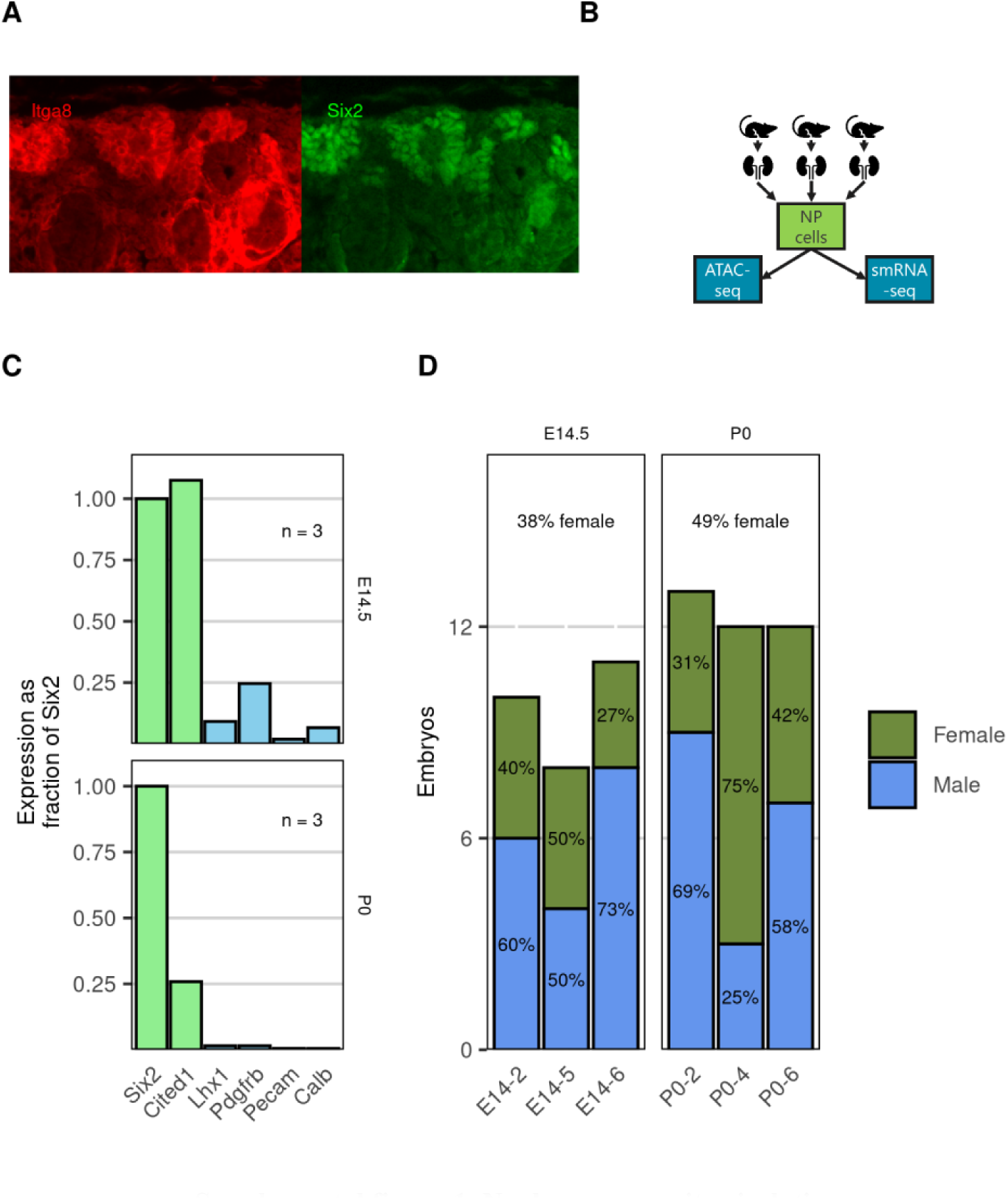
Nephron progenitor isolation. **A)** Immunofluorescence shows co-expression of *Six2* and *Itgα8* surface markers in developing mouse NPs. **B)** Nephron progenitors were isolated from pooled kidneys collected from the same litter at E14.5 and P0. **C)** Quantitative PCR (qPCR) was used to confirm enrichment of NP-specific markers *Six2* and *Cited1* relative to markers for ureteric bud (*Calb*), renal stroma (*Pdgfrb*), endothelial (*Pecam*), and renal vesicle / developing renal structure (*Lhx1*) cell types. **D)** The sex of each embryo was determined by PCR and tallied for each sample pool.

**Supplemental figure 2.**
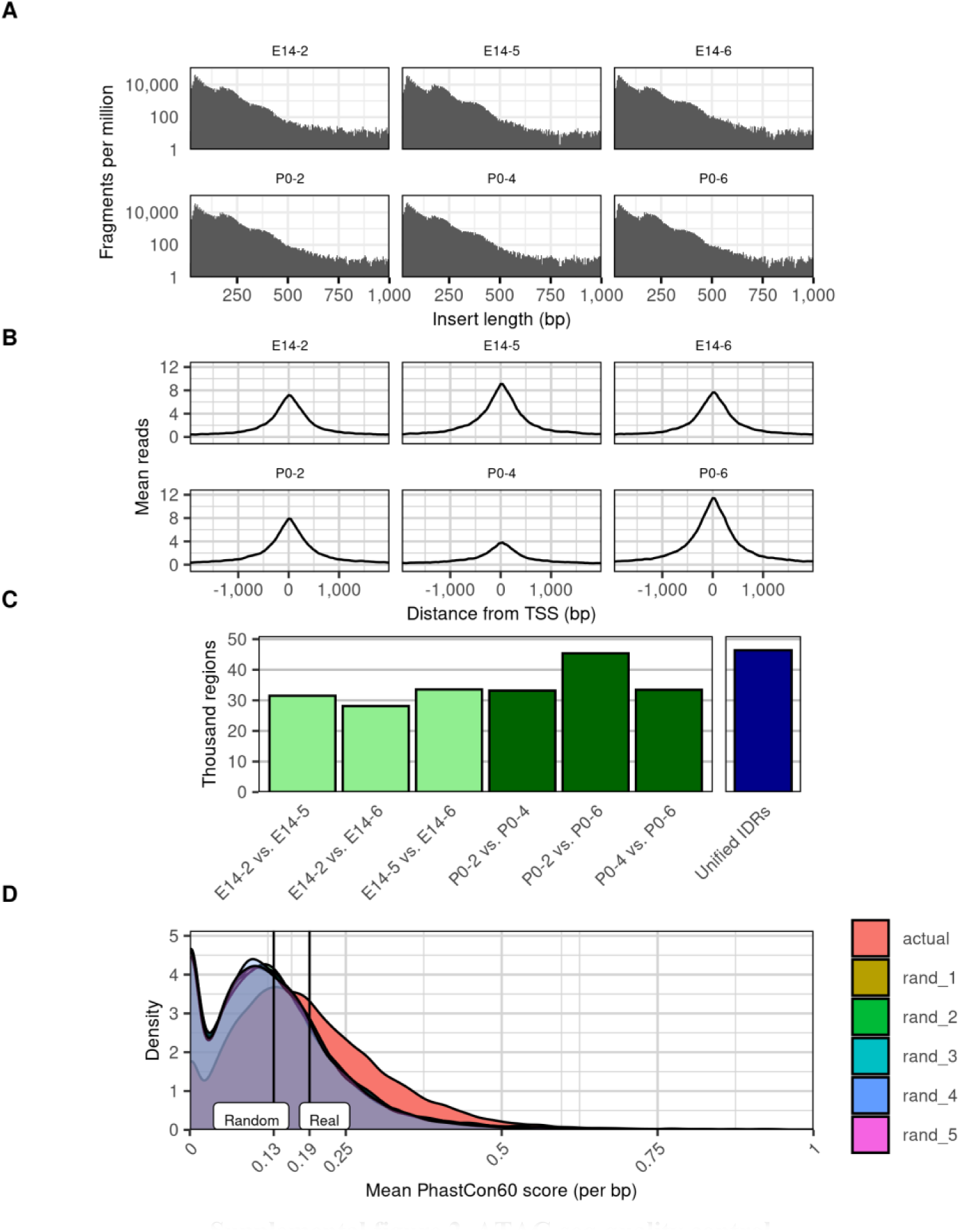
ATAC-seq quality control. **A)** The frequency of read lengths resulting from ATAC-seq exhibit periodicity corresponding to sub-nucleosomal, mono-nucleosomal, and di-nucleosomal distances. **B)** Tn5 insertion events are enriched at transcription start sites (TSSs). **C)** Of all peaks in ATAC-seq signal identified by MACS2, only those which are consistent between at least one pair of replicates using the irreproducible discovery rate (IDR; FDR = 0.1) are considered. The total number of IDR peaks identified in pairwise comparisons of peak sets from each sample are shown in green, and the union of all such IDR peaks is shown in dark blue. **D)** Distribution of average PhastCon60 scores per basepair for all IDRs (orange) compared to five random relocations of the IDR peaks in the genome.

**Supplemental figure 3.**
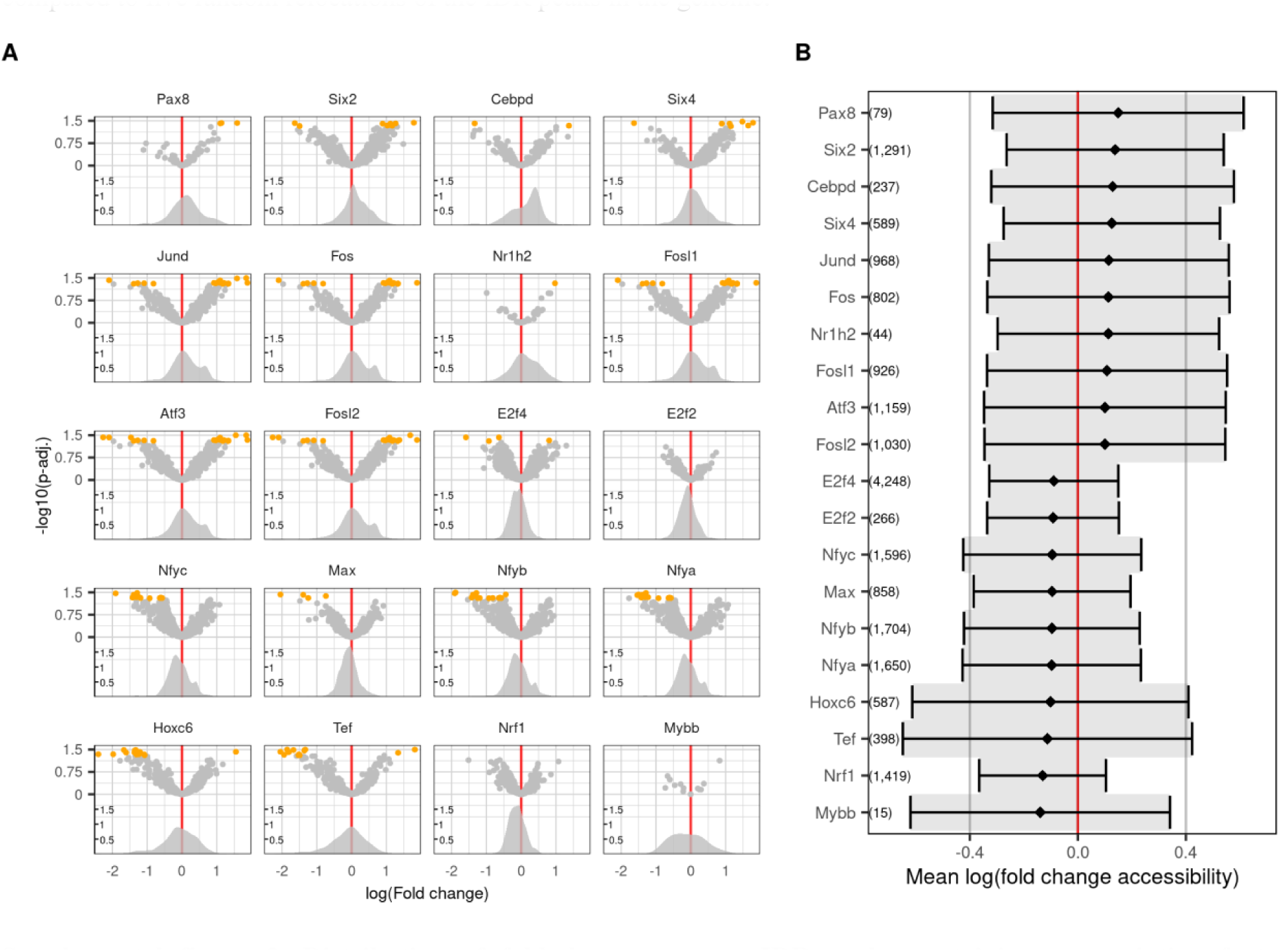
Distribution of fold-changes among IDR peaks containing transcription factor footprints. **A)** IDRs containing at least one transcription factor footprint for DNA binding proteins are plotted as a volcano plot in the top portion of each facet, with log2 fold change on the x-axis and -log10 of the peak’s adjusted P-value on the y axis. DARs are highlighted in orange, and the overall distribution of IDR fold changes are shown in the lower portion of each facet. **B)** Average log fold changes of IDR peaks containing one or more of the associated footprints. The top 10 greatest fold change increases and decreases are pictured, and the number of footprints averaged are shown in parenthesis.

**Supplemental figure 4.**
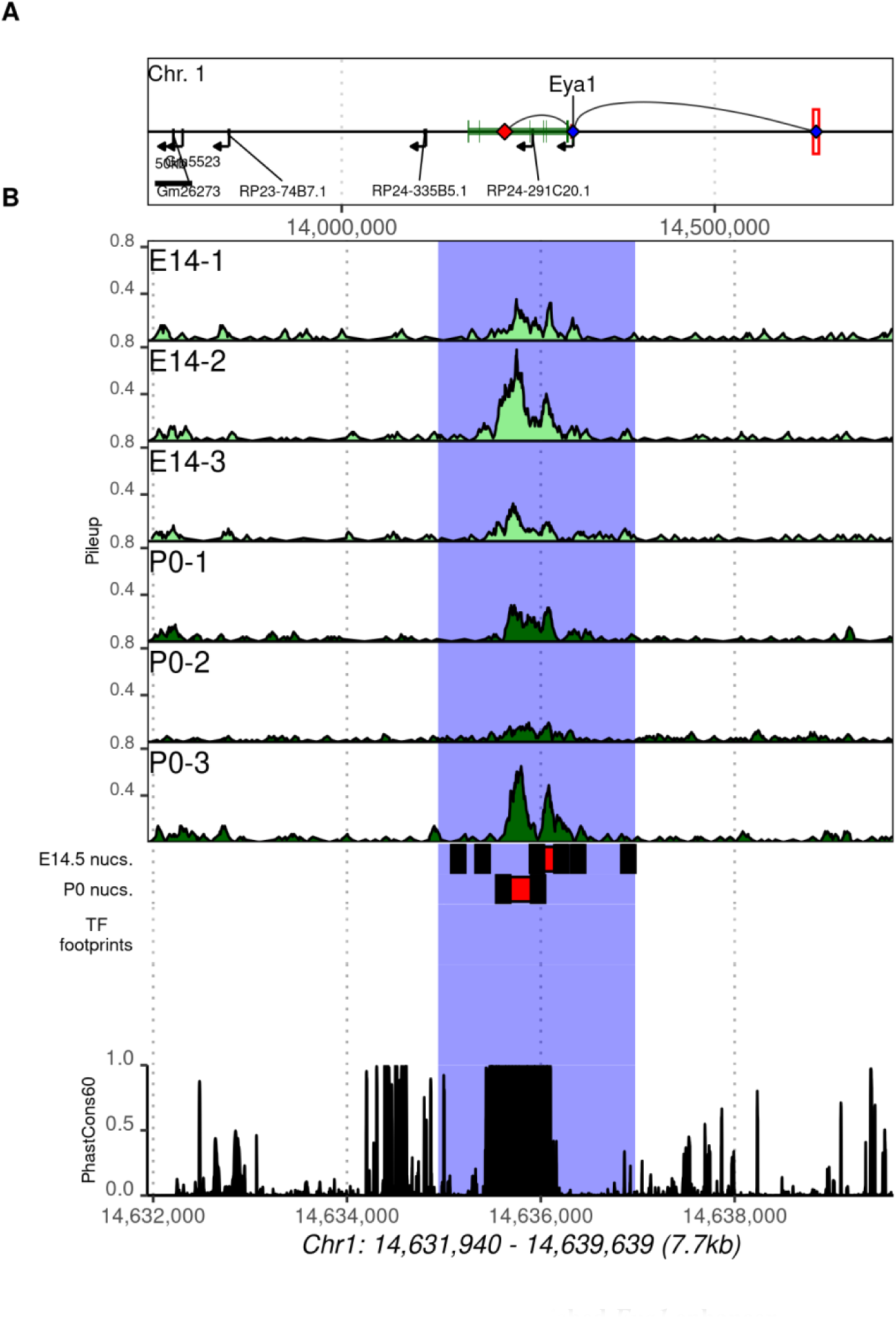
Published *Eya1* enhancer. **A)** Genomic location of enhancer of the *Eya1* transcript observed by Park et. al.(Park et al., 2012). **B)** ATAC-seq data from the regions surrounding the published enhancer, including (from top to bottom) E14.5 and P0 accessibility data (light green and dark green, respectively), nucleosomal positions (black rectangles) and nucleosome-free regions red rectangles), and sequence conservation among mammals. The detected IDR is highlighted in blue, and does not significantly change over time.

**Supplemental figure 5.**
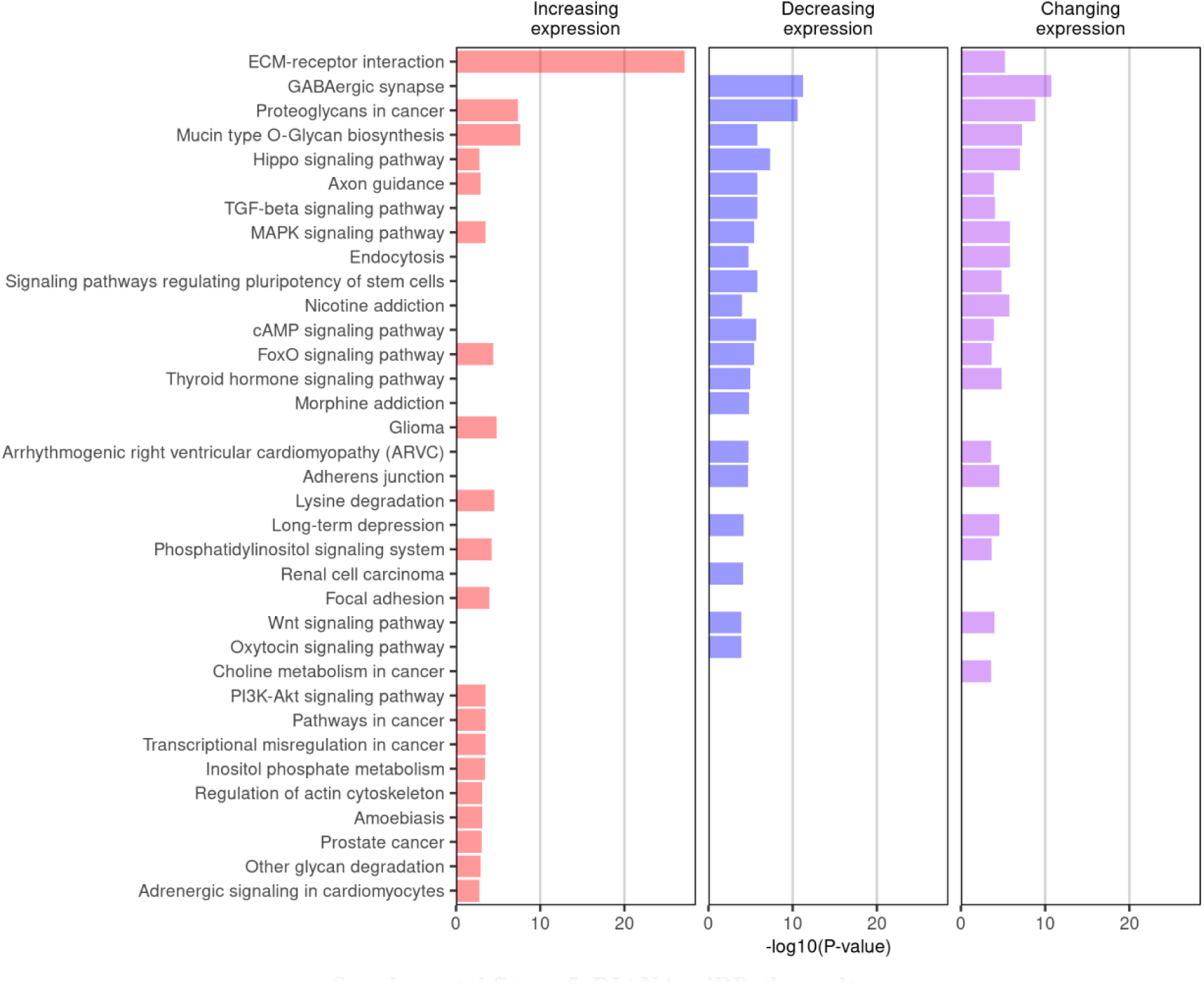
DIANA miRPath results. DIANA miRPath results for sets of increasing, decreasing, and changing (increasing and decreasing) miRNA.

